# Efficient identification of antimalarials with novel mechanism of action via multi-sample transcriptomic profiling

**DOI:** 10.1101/2025.02.27.640686

**Authors:** Ryuta Ishii, Takaya Sakura, Jing Hong, Kazuya Yasuo, Kazunari Hattori, Paul A Willis, Teruhisa Kato, Daniel Ken Inaoka

## Abstract

Malaria poses a significant health burden to endemic countries, necessitating the development of new antimalarials with novel mechanisms of action to combat emerging drug resistance. However, target identification methods for antimalarial compounds selected through phenotypic screening are currently limited, which has been a major obstacle in the field of antimalarial drug discovery. Here, we aim to address this need using perturbation transcriptomics to efficiently select antimalarial compounds targeting new pathways or molecules. The method involves comparing the transcriptomic changes by drug perturbation in the asexual blood stage of the malaria parasite *Plasmodium falciparum.* Using this method, we identified several compounds that exhibited distinct transcriptomic profiling from a diverse set of antimalarials with known modes of action, indicating their potential to target different pathways. In addition, we successfully identified a novel histone deacetylase inhibitor using this approach and chemogenetically validated its target through *in vitro* evolution. These results highlight the effectiveness of this platform in accelerating antimalarial drug discovery.

## Introduction

Malaria is a devastating disease caused by *Plasmodium* parasites transmitted to humans through the bites of *Anopheles* mosquitoes. According to the World Health Organization, there were an estimated 263 million malaria cases and 597 thousand deaths in 2023^1^. Particularly in sub-Saharan Africa, where 94% of cases and 95% of deaths occur, malaria poses significant public health challenges, leading to economic and developmental issues despite the availability of several treatment options.

The first-line treatment of uncomplicated malaria since 2006 has been artemisinin-based combination therapies (ACT), the combination of artemisinin derivatives and different class partner drugs. Over 200 million ACT treatment courses are used yearly^1^, and the overall treatment success rate is higher than 90%^2^, however, emerging artemisinin partial resistance is confirmed in many countries, including in sub-Saharan Africa^3–7^. In addition, there are emerging reports of potential resistance to the ACT partner drugs. One of the strategies to combat emerging resistance is developing a new class of antimalarials that acts by novel mechanisms of action (MoAs) and can replace artemisinin derivatives. A single oral dose regimen and rapid parasite clearance are essential to replace artemisinin derivatives and avoid the emergence of a new resistance^8^. Many groups have been working on fulfilling those criteria, however, no single-dose cure product has yet made it to market. Continuous drug discovery studies are required to reduce the burden of malaria toward elimination and eradication.

Although the target-based approach is a dominant methodology in drug discovery, a comprehensive analysis suggested its inefficiency^9^. That is also true in antimalarial drug discovery; despite the dedication of many groups for more than a decade, none of the target-based antimalarial candidates have succeeded yet^10^, and most of the new class pipeline drugs in clinical trials are developed from phenotypic screening^11–17^. Phenotypic screening may allow researchers to easily select potent compounds, however, the uncertainty of the MoAs can preclude quick drug development, mainly due to issues in safety profiles and structure-activity relationships^18^. Furthermore, without identifying the actual MoAs, there is always a risk of developing new compounds with the same MoA as an existing drug, which cannot then be used as a next-generation antimalarial combination therapy. Therefore, several target identification techniques have been developed for identifying targets of antimalarials, such as click-chemistry pull-down, cellular thermal shift assay, and multi-omics approach^19^. These techniques sometimes provide precise results; for example, the targets of plasmepsin inhibitor WM382 were identified by cellular thermal shift assay^20^. However, given that the targets of antimalarials in the drug development pipeline are not specified in many cases^16,21,22^, there are limitations of those techniques, depending on the targets or compounds, and a new approach to antimalarial target identification is required. It would be particularly advantageous if any new methodology could be applied to tens of compounds, allowing it to be used to prioritize the most promising hits emerging from high throughput screens. Many of the existing methods are low throughput or required significant resources and time, making them unsuitable for screening a large number of compounds in a timely manner.

RNA-seq analysis is a powerful technique that uses high-throughput sequencing to detect and quantify gene expression levels. It allows researchers to analyze a wide range of RNA populations, including mRNA and total RNA, providing valuable insights into biological processes. Transcriptome patterns of cells can be changed by any stimulation, including drug treatment, and can be a strong tool for drug discovery. It was reported that high-throughput RNA-seq analysis could distinguish MoAs of compounds by clustering based on transcriptomic changes of mammalian cell lines after drug perturbulation^23^. Identifying the exact MoA of a compound was also possible by using the same technology with comparing transcriptome patterns of known MoA compounds^24^, showing the usefulness of this technique in phenotypic drug discovery. RNA-seq analysis has also been used in research on the malaria parasite *Plasmodium falciparum* because its genome is one of the best annotated eukaryotic genomes^25^. It has been demonstrated that RNA-seq analysis can provide new insights into a variety of research, from *in vitro* single-cell analysis for identifying sexual differentiation drivers to the identification of transcriptional patterns in field isolates related to cerebral malaria^26,27^. However, multi-sample transcriptomics has never been applied to the asexual blood stage in the field of antimalarial drug discovery, Here, we propose a method to detect and compare transcriptomic changes of *P. falciparum* by drug perturbation, which could estimate the MoAs of antimalarial drugs and led to identification of a compound with a novel MoA. Our transcriptome clustering identified a novel *P. falciparum* histone deacetylase 1 (*Pf*HDAC1) inhibitor from previously uncharacterized phenotypic antimalarial hits, demonstrating the effectiveness of this method as a powerful tool for target identification studies.

## Results

### RNA-seq analysis of drug perturbation of *P. falciparum* asexual blood stage

To establish a method for efficiently detecting transcriptomic changes in *P. falciparum* after drug perturbation, we implemented a 96-well plate-based procedure. Since mRNA expression patterns significantly change throughout the different stages of the blood stage parasite^28^, we separately collected RNA from the tightly synchronized ring, trophozoite, and schizont stage parasites after treatment with each compound (Fig. 1). To comprehensively understand how the MoA of a compound affects transcriptomic changes, we tested 63 compounds in triplicate for this study (Table S1). Of these, 9 compounds were structurally new and identified from Shionogi & Co., Ltd. chemical library^29^, while others were compounds demonstrated to have antimalarial activity including a diverse set of compounds from Medicines for Malaria Venture (MMV) having either a known mode of action or compounds refractory to resistance where a mode of action could not be determined. RNA purification and reverse transcription were also performed in a 96-well plate, after which samples were collected for library preparation and Illumina sequencing. As a result, the number of read counts and detected genes varied according to the developmental stage, consistent with their RNA content^30^ (Fig. 2). Only four out of 576 samples were excluded during quality control based on a detected gene count 500 or below. We concluded that antimalarial compound-treated parasites had a sufficient amount of RNA and could be used for the RNA-seq analysis.

**Figure 1.**
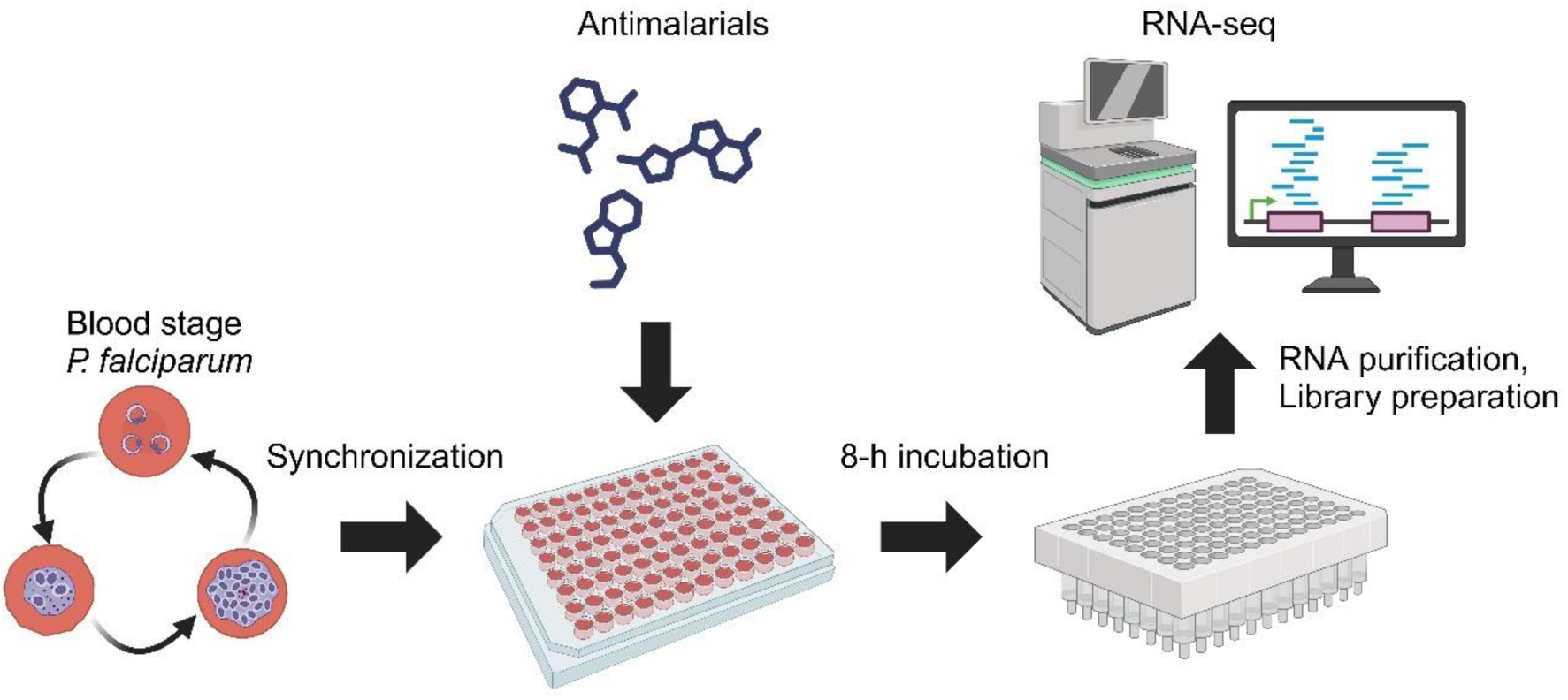
Overview of the assay procedure. *P. falciparum* was tightly synchronized to the ring, trophozoite, or schizont stage by sorbitol synchronization at various time points and dispensed onto compound plates. After an 8-hour incubation, total RNA from each sample was purified using a 96-well spin column, followed by cDNA library preparation and sequencing using a next-generation sequencer.

**Figure 2.**
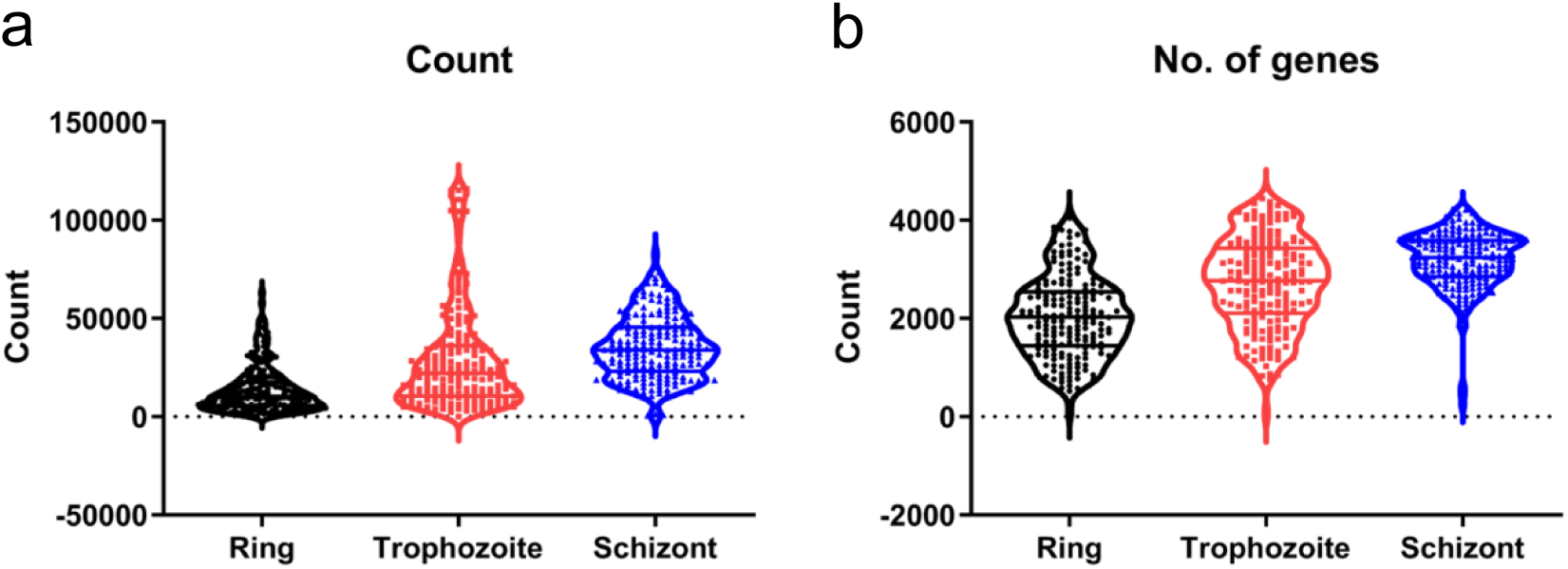
RNA-seq qualities of drug-treated parasites. Distribution of **a** read count and **b** number of genes detected by RNA-seq. Each parasite stage includes 192 samples from drug-treated asexual *P. falciparum* parasites.

### Transcriptomic changes can distinguish MoAs of antimalarials

For further analysis, we utilized 1000 highly variable genes to maximize the differences in expression patterns among each sample. We performed dimensionality reduction using Uniform Manifold Approximation and Projection (UMAP)^31^, followed by clustering analysis. This analysis revealed several clusters of samples that exhibited distinct changes in mRNA expression (Fig. 3). The clustering results for the ring stage suggested that this stage may not be suitable for observing MoA-based transcriptomic changes, as it only showed clusters with a wide range of different MoAs. However, the results for the trophozoite and schizont stages revealed more clusters, some of which included a small number of compounds that may have the same MoA (Table 1). For example, Cluster No. 6 in the schizont stage includes cytochrome *bc_1_* inhibitors and the dihydroorotate dehydrogenase (DHODH) inhibitor DSM265, all of which should lead to the same phenotype according to the previous findings showing that these compounds lose their antimalarial activities when yeast DHODH is overexpressed in the blood stage of the parasite^32,33^. Furthermore, MMV021735 has been reported to have metabolic profiles similar to those of DSM265 and atovaquone^34^, indicating that our RNA-seq analysis has the potential to detect inhibition of the mitochondrial pathway by this compound as well. Overall, the schizont stage showed more clusters with distinct MoAs: Cluster No.3 contains hemozoin formation inhibitors (chloroquine^35^, ferroquine^36^), Cluster No. 4 contains translation inhibitors (M5717^37^, MMV670325^38^, MMV674143^39^, and MMV009952^40^), Cluster No. 5 contains dihydrofolate reductase–thymidylate synthase (DHFR-TS) inhibitors (pyrimethamine^41^ and MMV027634^42^). Our phenotypic hit with a novel chemotype was also included in Cluster No. 3, potentially indicating the finding of a new class of hemozoin formation inhibitor. In addition, the result of Cluster No. 10 in the trophozoite stage suggests that MMV011438 may function as a new inhibitor of acetyl-coenzyme A synthetase (AcAS), despite the presence of a resistance gene that is essential for activating the compound^43^. Further investigation into the MoAs of these novel compounds is required.

**Figure 3.**
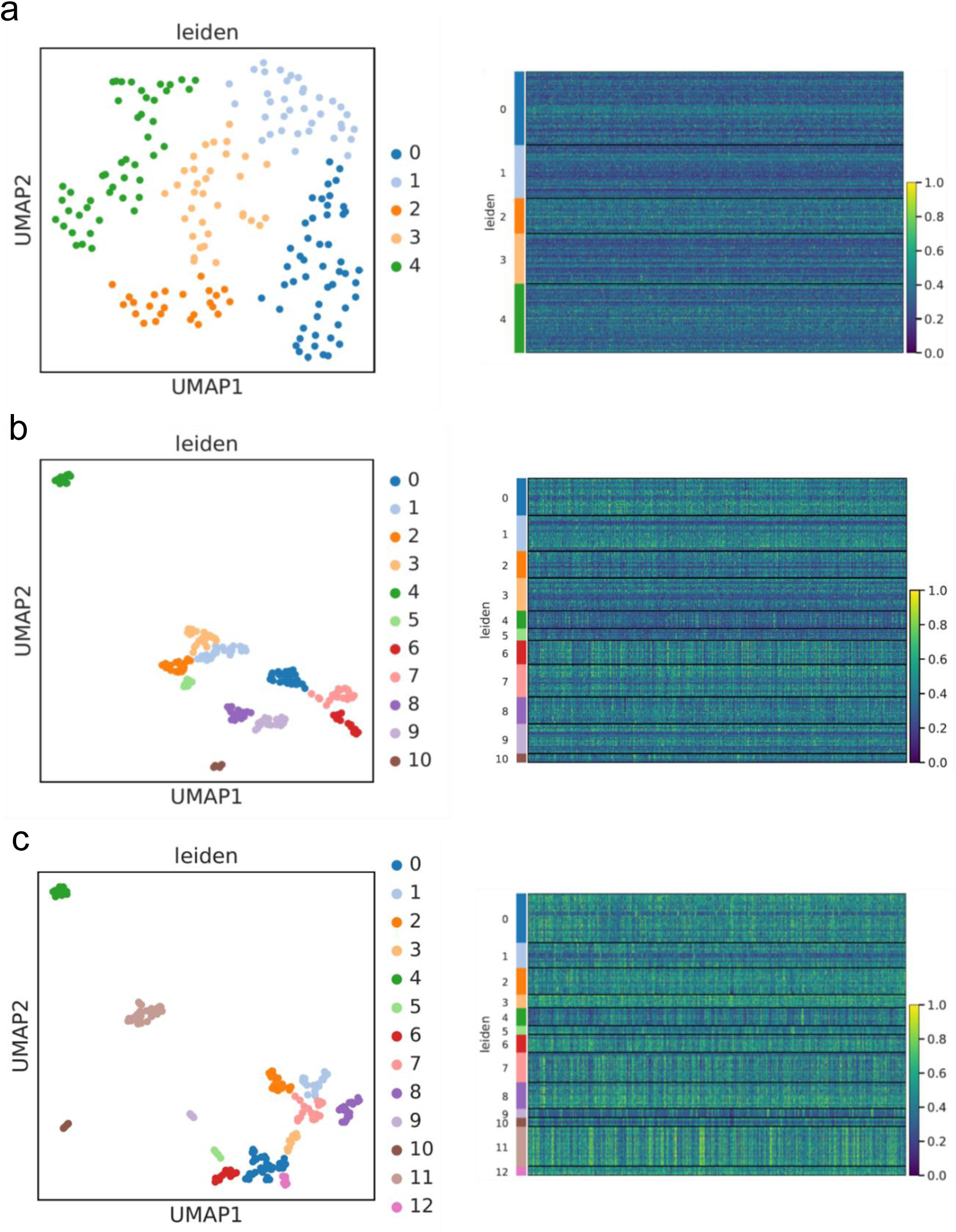
Clustering analysis based on transcriptomic changes of parasites after drug treatment. The results of mapping and clustering of transcriptomes from **a** the ring stage, **b** the trophozoite stage, and **c** the schizont stage of *P. falciparum*. The Uniform Manifold Approximation and Projection (UMAP) of quality-filtered transcriptome (left panels). Cluster numbers for each stage are indicated by different colors. Heatmaps of normalized gene expression levels displayed according to the clusters (right panels). Clusters with distinctive MoAs are summarized in Table 1. The number of samples for the ring, trophozoite, and schizont stages were 191, 191, and 190, respectively.

**Table 1.**
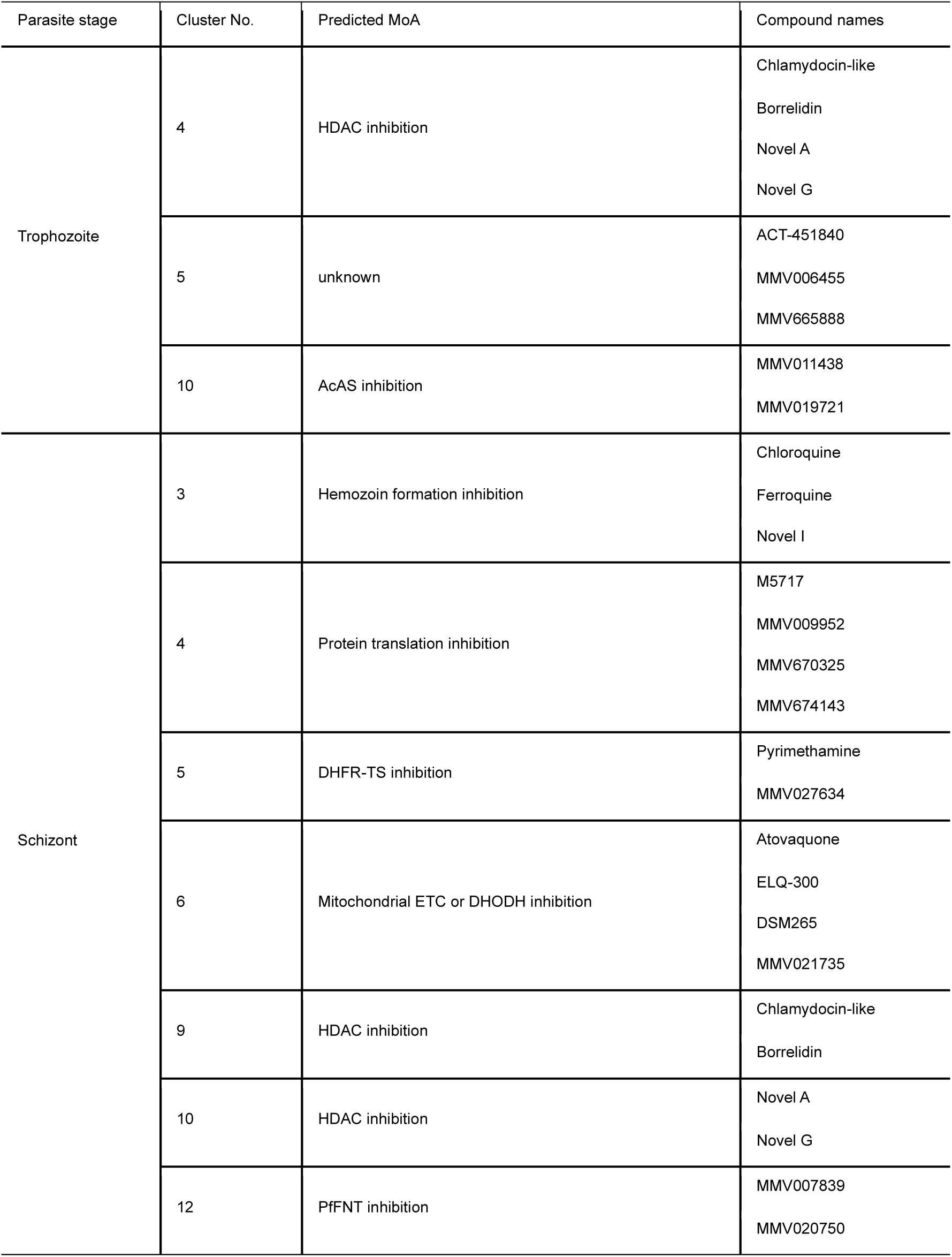
List of compounds in distinctive clusters. Clusters of compounds with similar or small numbers of MoAs are listed. All the triplicate samples in the compounds listed above were in the same cluster. MoA: mechanism of action, HDAC: histone deacetylase, AcAS: acetyl-CoA synthetase, DHFR-TS: bifunctional dihydrofolate reductase-thymidylate synthase, ETC: electron transport chain, DHODH: dihydroorotate dehydrogenase, PfFNT: *P. falciparum* formate-nitrite transporter.

### RNA-seq analysis identified a novel *Pf*HDAC1 inhibitor

The results of Cluster No. 4 in the trophozoite stage were inconclusive because it included compounds with two different MoAs: chlamydocin-like HDAC1 inhibitor^44,45^ and threonine-tRNA synthetase (TRS) inhibitor Borrelidin^46^. To determine the MoA of the phenotypic hits, we conducted Novel A compound resistance selection using the Dd2 strain. After 23 days of single-step selection at a concentration equal to 5-fold EC_50_, viable parasites were observed, followed by cloning, which resulted in the obtaining of five resistant clones. All the clones exhibited similar fold changes to Novel A (Fig. S1), and whole genome sequencing identified single nucleotide variants (SNVs) of three genes, HDAC1 (PfDd2_090030700), Cyclic nucleotide-binding domain containing protein (PfDd2_140022100) and conserved *Plasmodium* protein, unknown function (PfDd2_140047100) as common SNVs in all clones (Fig. S2). According to the gene conservation among *P. falciparum* strains and essentiality, we focused on the L137F mutation in PfHDAC1. To further validate the effect of the L137F mutation in Novel A resistance, we generated the Dd2 strain with or without the PfHDAC1 L137F mutation (Dd2-L137F) using CRISPR-Cas9 mutagenesis (Fig. 4a). Drug selection, followed by cloning, allowed us to obtain three clones each with confirmed synthetic DNA insertions through diagnostic PCR (Fig. S3). Clones with the L137F mutation demonstrated EC_50_ fold changes similar to the Novel A resistant strains (Fig. S4), confirming that the L137F mutation is responsible for the resistance in these strains. We further corroborated the effect of the L137F mutation on resistance to PfHDAC1 inhibitors using molecular docking with a Novel A derivative Compound 1, which also showed lower activity against the L137F mutant strains (Figs. 4b, 4c). In this analysis, the bulky moiety of the phenylalanine mutant obstructed the binding of Compound 1 into the active site of PfHDAC1 (Fig. 4d). Based on these findings, we concluded that the Novel A compound, and presumably the structurally similar Novel G compound, are a new-class of PfHDAC1 inhibitors, underscoring the utility of our RNA-seq approach in identifying the MoAs of novel antimalarials.

**Figure 4.**
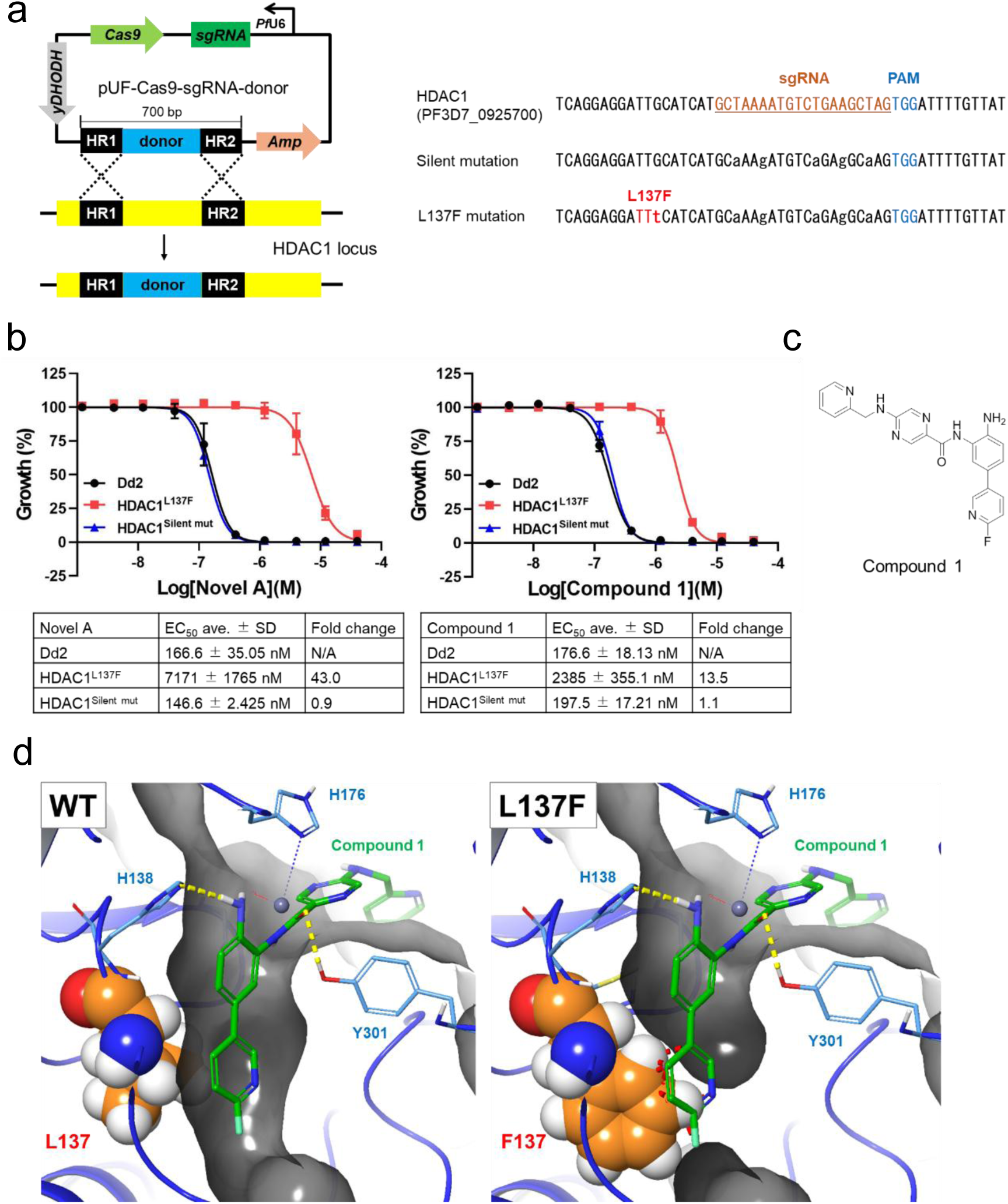
A novel compound series targeting PfHDAC1 and their activity loss due to L137F mutation in PfHDAC1. **a** Schematic representation of donor DNA insertion into the HDAC1 locus using CRISPR-Cas9 (left panel). Five silent mutations are inserted into the sgRNA target sequence of both control and L137F mutation donor DNAs (right panel). **b** Dose-response curves of Novel A (left panel) and Novel A derivative Compound (right panel) for the parental Dd2 line, HDAC1:L137F mutant lines, and HDAC1:silent mutation control lines (four technical replicates and three independent experiments, mean ± SD). **c** The structure of Compound 1. **d** PfHDAC1/ Compound 1 complex model for the wildtype (left panel) and L137F mutant (right panel). Compound 1 is depicted in green, and the L137 or F137 amino acid is highlighted with the electron density of atoms. Yellow dot lines represent the hydrogen bonds between the compound and amino acids. The red dot line indicates the collision between F137 and Compound 1. yDHODH: yeast dihydroorotate dehydrogenase, sgRNA: single guide RNA, HR: homologous region, Amp: ampicillin-resistant gene, PAM: protospacer adjacent motif.

## Discussion

In the present study, we utilized multi-sample RNA-seq to estimate the MoAs of a wide variety of antimalarials in *P. falciparum* for the first time. This method is time-efficient, only requiring an 8-hour drug treatment of the parasite in a 96-well plate. Furthermore, recent advancements in next-generation sequencers have made it cost-effective and capable of processing larger sample sizes. Although the RNA content of the parasite at different developmental stages affected read counts and the number of detected genes, antimalarial drug treatment was compatible with performing the RNA-seq analysis and subsequent clustering based on changes in transcriptomic patterns. We evaluated 63 compounds with at least 35 different MoAs, resulting in 4, 10, and 12 clusters in the ring, trophozoite, and schizont stages, respectively. These findings suggest limitations in our method, as not all antimalarial treatments induce transcriptomic changes in parasites, and it is unpredictable which compounds will form distinctive clusters until tested. In addition, we did not observe MoA-specific clusters in the ring stage, potentially indicating that our method is unsuitable for assessing MoAs of ring stage-specific antimalarials. However, ferroquine, which is expected to be effective specifically in the ring stage^34^, was detected as one of the hemozoin formation inhibitors in the trophozoite stage, suggesting that ring stage-specific inhibitors may also affect the transcriptome of parasites in different stages. Further investigation is required to fully understand the behavior of ring stage-specific antimalarials in our method.

It is worth noting that compounds targeting different proteins but predicted to have the same phenotypic outcomes clustered together, such as translation inhibitors and electron transport chain inhibitors. This demonstrates that our method can detect the transcriptomic response of parasites to compounds and can be a powerful tool for identifying antimalarials with novel MoAs. In the trophozoite stage, Cluster No. 5 includes only three antimalarials with unknown MoAs, suggesting that they may share a novel target. Of these three compounds, MMV1542945, also known as ACT-451840, only exhibited resistance during the minimum inoculum for resistance (MIR) studies^47^. All resistant strains had mutations in *P. falciparum* multidrug resistance-1 (PfMDR1), but there is no clear evidence of a direct interaction between PfMDR1 and ACT-451840. Assessing the efficacy of MMV006455 and MMV665888 against PfMDR1 mutants would provide new insights into understanding the MoA of ACT-451840. Another example of compounds with unknown MoAs that exhibited distinguishable transcriptomic patterns is Cluster No. 12 in the schizont stage, comprising MMV007839 and MMV020750. It has been demonstrated that MMV007839 inhibits *P. falciparum* formate-nitrite transporter (PfFNT)^48,49^, while MMV020750 may be a weak aminopeptidase inhibitor or β-hematin inhibitor^50,51^. Given that the relations between reported activities of MMV020750 and PfFNT functions are unclear, MMV020750 may directly inhibit PfFNT. Conducting a PfFNT functional assay would be necessary to confirm this MoA.

*In vitro* resistance selection and CRISPR-Cas9 mutagenesis results demonstrated that Cluster No. 4 in the trophozoite stage consists of HDAC inhibitors, while Borrelidin is reported to be a TRS inhibitor. Borrelidin has been found to have different functions in other systems, such as angiogenesis inhibition and apoptosis induction^52,53^, and it is also known as a cyclin-dependent kinase (CDK) inhibitor of *Saccharomyces cerevisiae*^54^. CDKs are conserved in *P. falciparum* and expressed in the asexual blood stage, where they are believed to have diverse functions^55^. Among these CDKs, *P. falciparum* CDK-related kinase 3 (Pfcrk-3) was found to be associated with protein complexes exhibiting HDAC activity and is essential for parasite proliferation^56^. This indicates that inhibition of Pfcrk-3 by Borrelidin may have resulted in the inhibition of HDAC and incorporation of Borrelidin into Cluster No. 4 in the trophozoite stage. The segregation of HDAC inhibitors into Clusters No. 9 and No. 10 in the schizont stage suggests they may have slightly different functions, possibly due to their specificities against other HDACs expressed in the asexual blood stage^57^. However, *in vitro* functional assays will be required to investigate further details.

In conclusion, we have developed a method for conducting RNA-seq of the blood stage malaria parasite *P. falciparum* after antimalarial treatment using a 96-well plate. Clustering based on transcriptomic changes in the drug-treated parasite successfully predicted the MoAs of several antimalarials. This assay is believed to be advantageous in identifying antimalarials with novel MoAs, thereby accelerating the discovery of next generation antimalarial drugs.

## Methods

### Parasite cultures

*Plasmodium falciparum* 3D7 and Dd2 parasites were maintained in RPMI1640 medium supplemented with 25 mg/L gentamicin, 50 mg/L hypoxanthine, 23.8 mM sodium bicarbonate, and 0.5% (w/v) Albumax II (Gibco). The culture was maintained at 2% hematocrit using O+ human erythrocytes obtained from The Japanese Red Cross Society. The parasite culture was maintained in the environment of mixed gas (5% O_2_, 5% CO_2_, 90% N_2_) at 37 °C. Parasitemia was determined using Giemsa staining or an XN-30 automated hematology analyzer (Sysmex)^58^.

### RNA-seq of drug-treated parasites

The *P. falciparum* 3D7 culture was sorbitol synchronized^59^ 72 h and 36 h before magnetic purification^60^ as a pre-synchronization step. The purified parasite was then added to new erythrocytes and allowed to invade for 3 h, followed by sorbitol synchronization to obtain a tightly synchronized parasite culture. After 8 h, 24 h, or 36 h from sorbitol synchronization for the ring, trophozoite, or schizont samples, respectively, 200 µL/well of the parasite culture was dispensed into 96-well plates with pre-dispensed compounds (200 nL/well of 10 mM DMSO solution using Echo 555 (Beckman Coulter)). The plates were incubated at 37 °C for 8 h and the total RNA from each sample was purified using NucleoSpin 96 (Takara) following the manufacturer’s instructions and stored at −80 °C. The cDNA library preparation from the purified RNA was performed using QIAseq UPX 3’ Transcriptome Kits (QIAGEN) following the manufacturer’s protocol, and Novogene performed next-generation sequencing.

### *In vitro* selection of Novel A resistant strains

The Dd2 strain was cultured with 1.15 µM (5-fold EC_50_) Novel A, with daily medium changes, for 5 days. After parasite clearance, parasitemia count and medium changes were performed three times a week until parasite regrowth and parasitemia reached above 1%. The resistant strain was cloned through limiting dilution of the obtained parasite culture. The growth inhibition of Novel A against ring-stage synchronized parasites was measured using the diaphorase-nitro blue tetrazolium assay, following previously described methods^29^.

### Whole genome sequencing of HDAC1 resistance mutants

Genomic DNA of Novel A resistant strains was extracted using Qiagen Blood and Tissue Kit (Qiagen). Illumina library preparation and sequencing were performed in Novogene. The fastq files of whole genome sequencing were created and aligned to reference genome of Dd2 (PlasmoDB-64_PfalciparumDd2_Genome, which was downloaded from https://plasmodb.org/common/downloads/release-64/PfalciparumDd2/fasta/data/) using BWA-MEM^61^ (v 0.7.18). SAMtools2 (v 1.18) was used for generating format converted files. Duplicate reads were removed by Picard tools MarkDuplicates (v 3.1.0), following with calling of single nucleotide variants (SNVs) which was performed using GATK HaplotypeCaller (v 4.4.0). The variants were narrowed down by removing the common variants of parent Dd2 strain and the resistant strains as well as filtered with QUAL > 1000, QD > 1.5, MQ > 30, DP> 5 using BCFtools^62^ (v 1.18). The common variants of five resistant strains were highlighted and annotated by SnpEff^63^ (v 5.1f) with genome annotation reference (PlasmoDB-64_PfalciparumDd2_AnnotatedCDSs). The variants that were evaluated as either HIGH or MODERATE were output. Regions of interest were inspected with Integrated Genome Viewer (v 2.16.2)^64^.

### Generation of HDAC1 resistance mutants

*P. falciparum* Dd2 strain and pUF-Cas9-pre-sgRNA^65^ obtained from Addgene were used for CRISPR-Cas9 genome editing. The sgRNA (Fasmac) was inserted into the *BtgZI* site of the plasmid. The synthesized donor DNAs purchased from Genewiz were amplified with primers containing 19 bp flanking plasmid homology sequences and inserted into the *NcoI* site using SLiCE^66^. The plasmid constructs were confirmed by Sanger sequencing. The plasmid was transfected into the parasite using a Gene Pulser Xcell (Bio-Rad). Fifty µg of plasmid was mixed with 150 µL of packed erythrocytes and brought to a final volume of 400 µL with cytomix (120 mM KCl, 0.2 mM CaCl_2_, 2 mM EGTA, 10 mM MgCl_2_, 25 mM HEPES, 5 mM K_2_HPO_4_, 5 mM KH_2_PO_4_; pH 7.6 adjusted with KOH)^67^. The transfected erythrocytes were mixed with magnetically purified Dd2 strain and cultured until the parasitemia reached 5%. The transgenic parasites were selected in a complete medium containing 1.5 µM DSM1 (Sigma-Aldrich) for 10 days and then maintained in a drug-free complete medium until parasite regrowth. Limiting dilution was performed to obtain transgenic clones, which were genotyped by diagnostic PCR and Sanger sequencing. The sgRNA, synthesized donor DNAs, and Sanger sequencing primers are shown in Supplementary Table S2.

### Generation of PfHDAC1/Compound 1 complex model

The homology model of PfHDAC1 was constructed following the method described in Mukherjee P., *et al.*^68^. Briefly, the structures of the templates, HDLP (PDB code: 1C3R) and HDAC8 (PDB code: 1T64), were obtained from the Protein Data Bank (PDB). The protein sequence of PfHDAC1 (Accession No. Q9XYC7) was retrieved from UniProt, and the amino acid sequences of 1C3R and 1T64 were aligned with the Q9XYC7 sequence. The alignment was performed using the Multiple Sequence Viewer application in Maestro from the Schrödinger suite^69^, with reference to the supplementary data in the paper. Subsequently, the homology model of PfHDAC1 was constructed using the Prime application. The molecular docking model was generated by docking to the homology model using Glide.

### Data analysis

The inhibition curves and EC_50_ values were determined using GraphPad Prism (v 8.4.3 GraphPad Software, Inc.). The raw data from the next-generation sequencer were processed, demultiplexed, and aligned to *P. falciparum* 3D7 annotated transcripts (PlasmoDB v. 64) using CLC Genomics Workbench (v 23.0.4, QIAGEN). Sample quality control, differential expression, and normalization were performed using Python (v 3.12.0). For data cleaning, genes with no expression in any sample were removed, and samples with detection of 500 or fewer genes were excluded. Read counts were log normalized, and 1000 highly variable genes were selected using the expression and dispersion for further analysis. The most significant principal components were employed for clustering, which was performed using Scanpy (v 1.10.0)^70^.

## Data availability

All sequence data that support the findings of this study have been deposited in NCBI sequence read archive (SRA) with the accession code PRJNA1226290.

## Code availability

The custom code scripts used to analyze data for this study are available at https://github.com/NEKKEN-MID/Transcriptome_MoA.

## Acknowledgements

The authors thank the Japanese Red Cross Society for providing human red blood cells and Dr. Ken-ichi Matsumura, Dr. Kenji Takaya, Ms. Rina Kaki, and Mr. Riku Ogasawara (Laboratory for Medicinal Chemistry Research, Shionogi & Co., Ltd.) for selecting Novel A-I compounds and synthesizing ELQ-300, M5717, MMV533, and ZY-19489. This work was supported by Shionogi & Co., Ltd., Osaka, Japan. Figure 1 was created in BioRender. Sakura, T. (2025) https://BioRender.com/q87j322

## Author contributions

R.I., T.S., and J.H. performed the experiments. R.I., T.S., J.H., and K.Y. analyzed the data. P.W. selected compounds. R.I., T.S., J.H., and D.K.I. wrote the paper. D.K.I., T.K., and K.H. directed the work. All authors directly participated in this work, including preparation of the manuscript and approval of the final version. All authors have read and agreed to the published version of the manuscript.

## Conflict of interest

R.I., K.Y., K.H., and T.K. are employees of Shionogi & Co., Ltd.

Nagasaki University and Shionogi & Co., Ltd., launched the Shionogi Global Infectious Diseases Division (SHINE) at the Institute of Tropical Medicine (NEKKEN), Nagasaki University. Under the collaboration agreement, Shionogi & Co., Ltd. provided the compounds pre-dispensed into the 96-well plates for the assay and contributed to discussions with the authors. Shionogi & Co., Ltd. did not influence the experimental design, data collection, data analysis, or interpretation.

## Supplementary information

**Figure S1.**
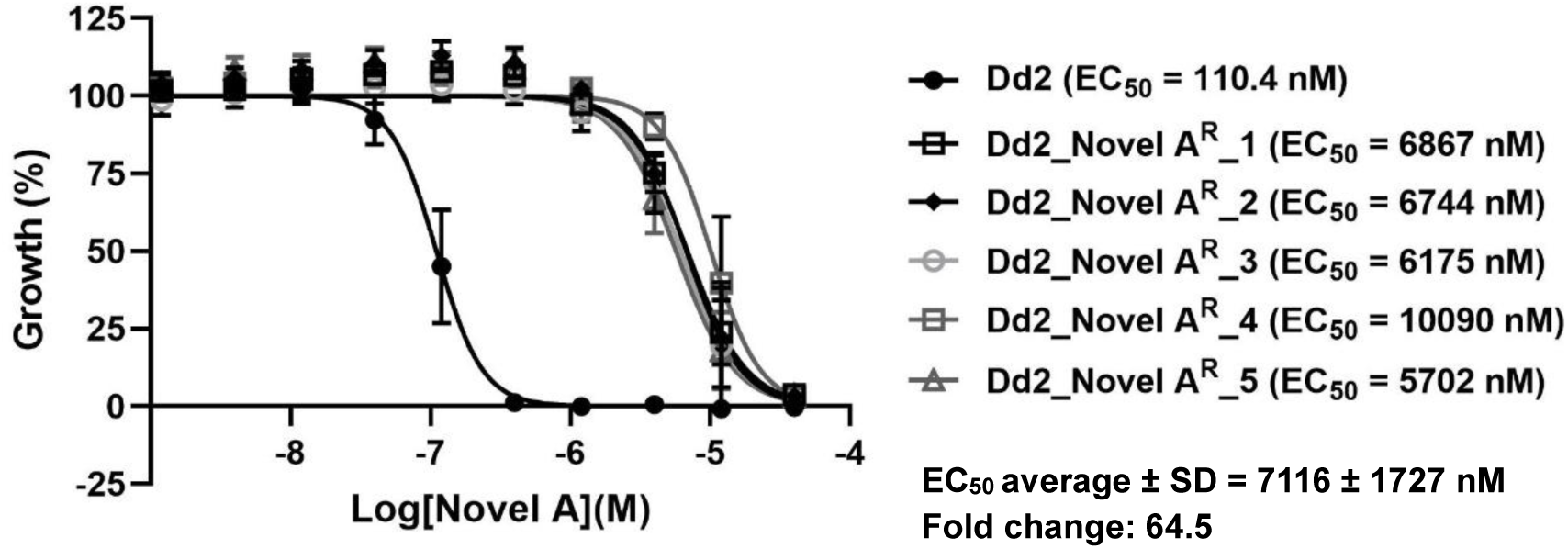
Efficacy of Novel A against obtained resistant strains. Dose-response curves of Novel A compound for the parental Dd2 line and five Novel A resistance clones (three technical replicates, mean ± SD).

**Figure S2.**
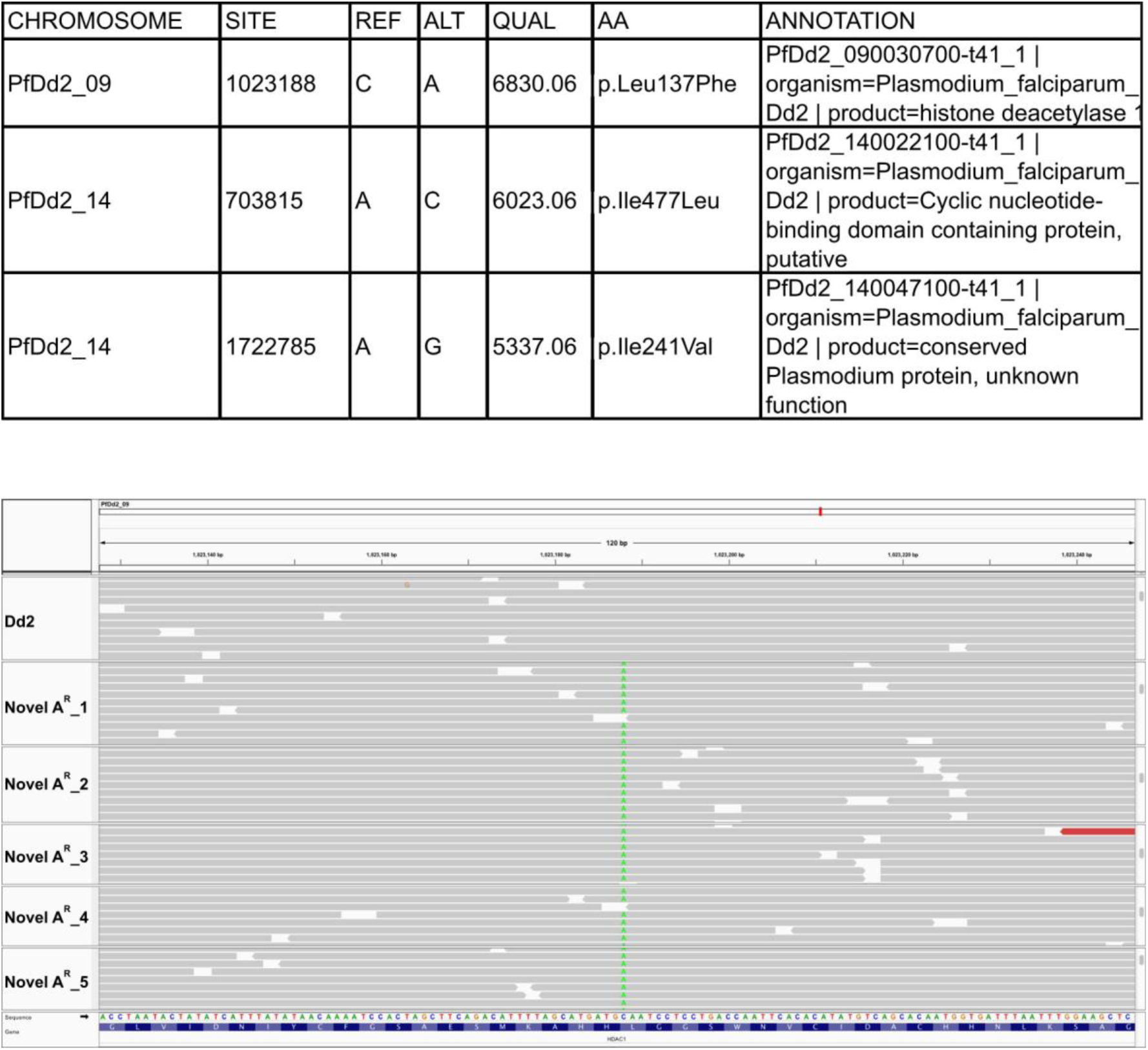
Whole genome sequencing of Novel A resistant strains. Three SNVs were identified among 5 resistant clones and L137F mutations of HDAC1 were visualized by Integrative Genomics Viewer (IGV).

**Figure S3.**
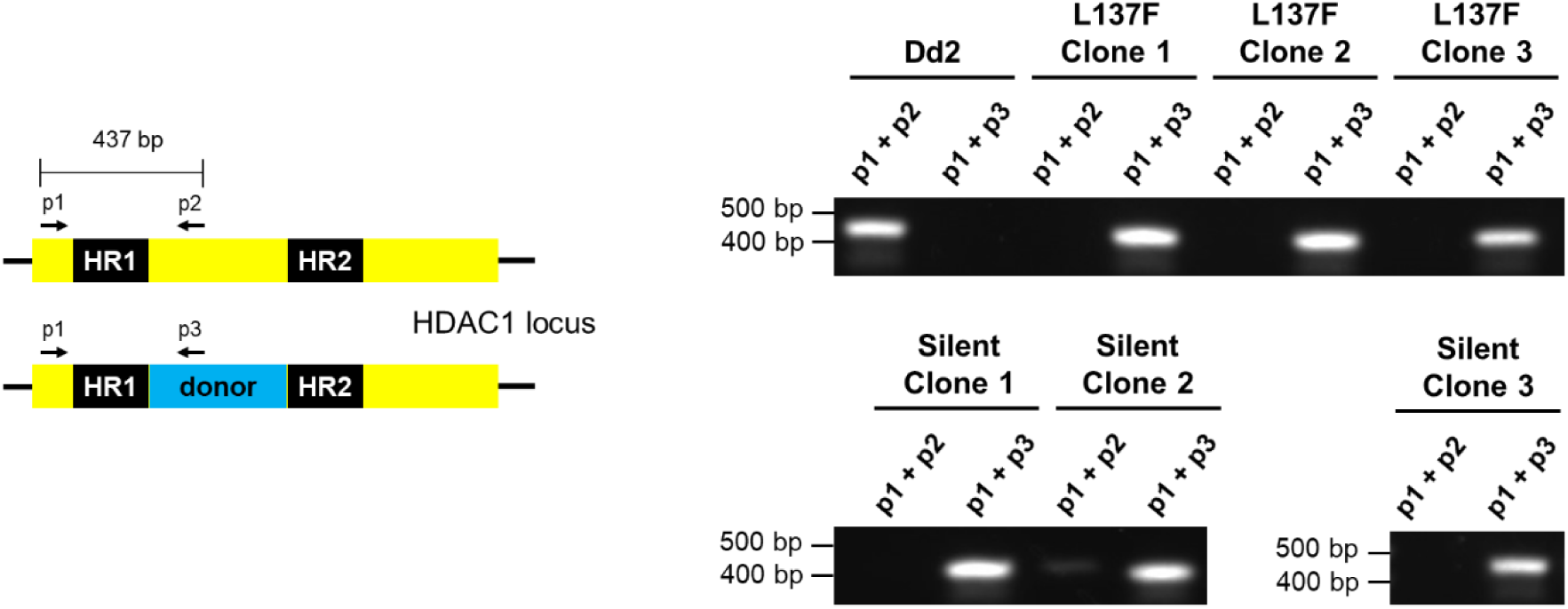
The result of diagnostic PCR to confirm insertion of donor DNA. Schematic representation of diagnostic PCR (left panel) and the results of diagnostic PCR for the parental Dd2 line and clones with donor DNA insertion (right panel).

**Figure S4.**
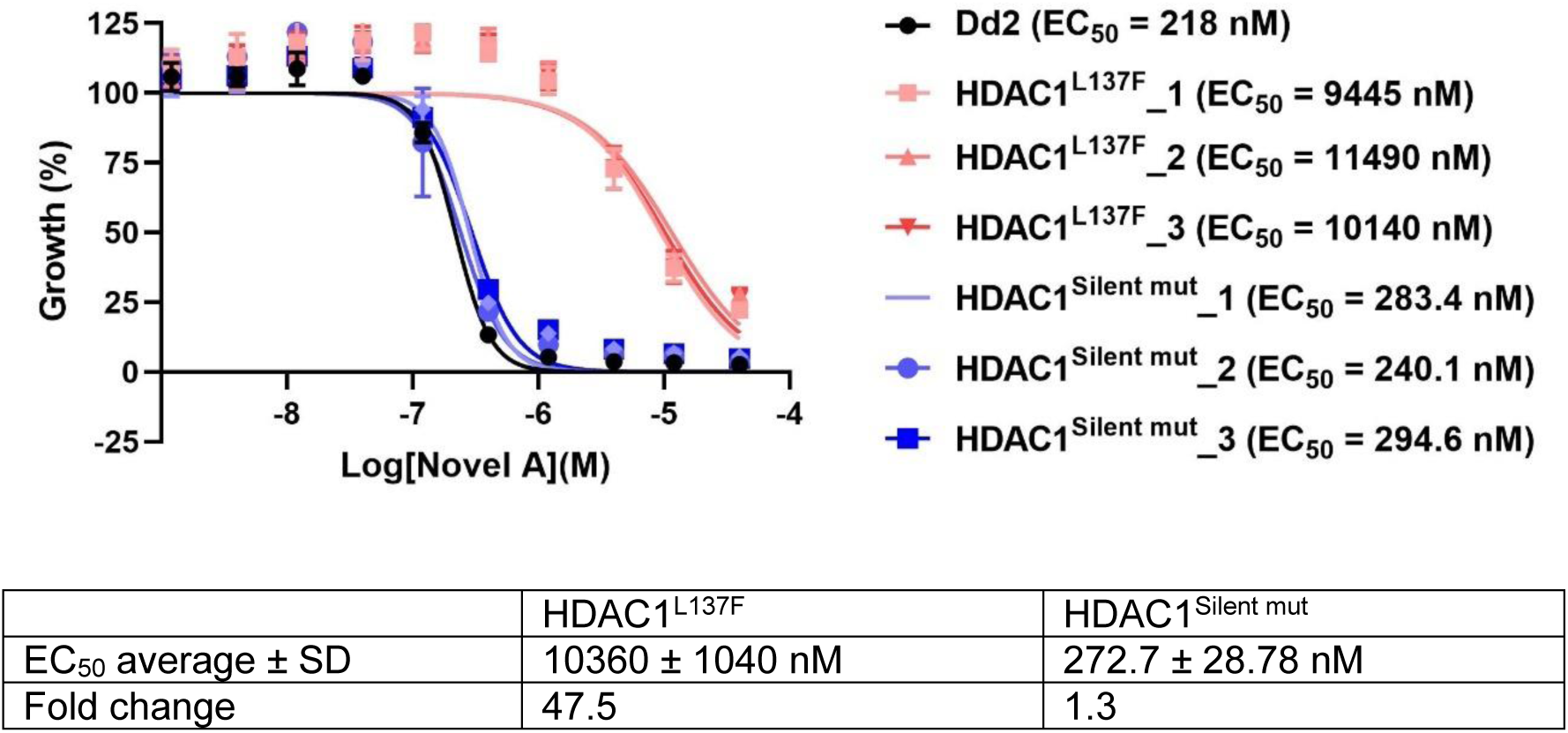
Efficacy of Novel A against transgenic L137F mutant strains. Dose-response curves of Novel A compound for the parental Dd2 line and five L137F mutant clones (three technical replicates, mean ± SD).

**Table S1.** List of tested compounds. (provided as an Excel file as well). The compounds with an MMV number were provided by MMV. Origins of other compounds are listed in Supplementary method. Chlamydocin-like compound is identical to molecule 3 of Traoré, M. D. M. *et al.*^45^, and Plasmepsin 2 inhibitor-like compound is identical to compound 8 of Hurley, K. A. *et al.*^71^, which is quite similar to plasmepsin 2 inhibitor^72^.

| Name | Expected target | Name | Expected target |
| --- | --- | --- | --- |
| MMV000041 (BCX4208) | purine nucleoside phosphorylase | Atovaquone | mitochondrial cytochrome bc1 complex (Qo site) |
| MMV006455 | unknown | BIX-01294 | DNA methyltransferase |
| MMV007224 | phospholipid-transporting ATPase 2 | Chlamydocin-like | histone deacetylase 1 |
| MMV007839 | formate-nitrite transporter | Chloroquine | hematin crystallization |
| MMV009952 (Halofuginone) | proline--tRNA ligase | Cipargamin | non-SERCA-type Ca2+ -transporting P-ATPase (PfATP4) |
| MMV011438 | unknown | Cytochalasin D | actin polymerization |
| MMV016838 | eukaryotic elongation factor 2 (predicted) | M5717 | eukaryotic elongation factor 2 |
| MMV019066 | protein farnesyltransferase subunit beta | Dihydroartemisinin | several proposed but not determined |
| MMV019313 | bifunctional farnesyl/geranylgeranyl diphosphate synthase | DSM265 | dihydroorotate dehydrogenase |
| MMV019721 | acetyl-CoA synthetase | ELQ-300 | mitochondrial cytochrome bc1 complex (Qi site) |
| MMV020438 | hexose transporter | Epoxomicin | proteasome |
| MMV020746 | ABC transporter I family member 1 | Ferroquine | hemozoin formation |
| MMV020750 | unknown | Ganaplacide | parasite internal protein secretory pathway but not fully understood |
| MMV021735 | unknown | INE963 | unknown |
| MMV022478 | unknown | Lumefantrine | unknown |
| MMV027634 | bifunctional dihydrofolate reductase-thymidylate synthase | Mefloquine | unknown |
| MMV029272 | dipeptidyl aminopeptidase 1 and ABC transporter I family member 1 | MMV048 | phosphatidylinositol 4-kinase |
| MMV084032 (Borrelidin) | threonine--tRNA ligase | MMV533 | unknown |
| MMV1542945 (ACT-451840) | unknown | Plasmepsin 2 inhibitor-like | plasmepsin 2 |
| MMV1580857 (Mupirocin) | isoleucine--tRNA ligase | Pyrimethamine | dihydrofolate reductase |
| MMV1792684 | cyclin-dependent-like kinase CLK3 | Quinine | unknown |
| MMV397264 (PA92) | non-SERCA-type Ca2+ -transporting P-ATPase (PfATP4) | ZY-19489 | unknown |
| MMV652007 (AN3661) | cleavage and polyadenylation specificity factor subunit 3 | Novel A | unknown |
| MMV665888 | unknown | Novel B | unknown |
| MMV665924 | acyl-CoA synthetase (ACS11) | Novel C | unknown |
| MMV665939 | unknown | Novel D | unknown |
| MMV666080 | unknown | Novel E | unknown |
| MMV667530 | 2-C-methyl-D-erythritol 4-phosphate cytidyltransferase (IspD) | Novel F | unknown |
| MMV668399 | amino acid transporter AAT1 and others | Novel G | unknown |
| MMV670325 (AN6426) | leucine--tRNA ligase | Novel H | unknown |
| MMV673964 | unknown | Novel I | unknown |
| MMV674143 (Cladosporin) | lysine--tRNA ligase |  |  |

**Table S2.**
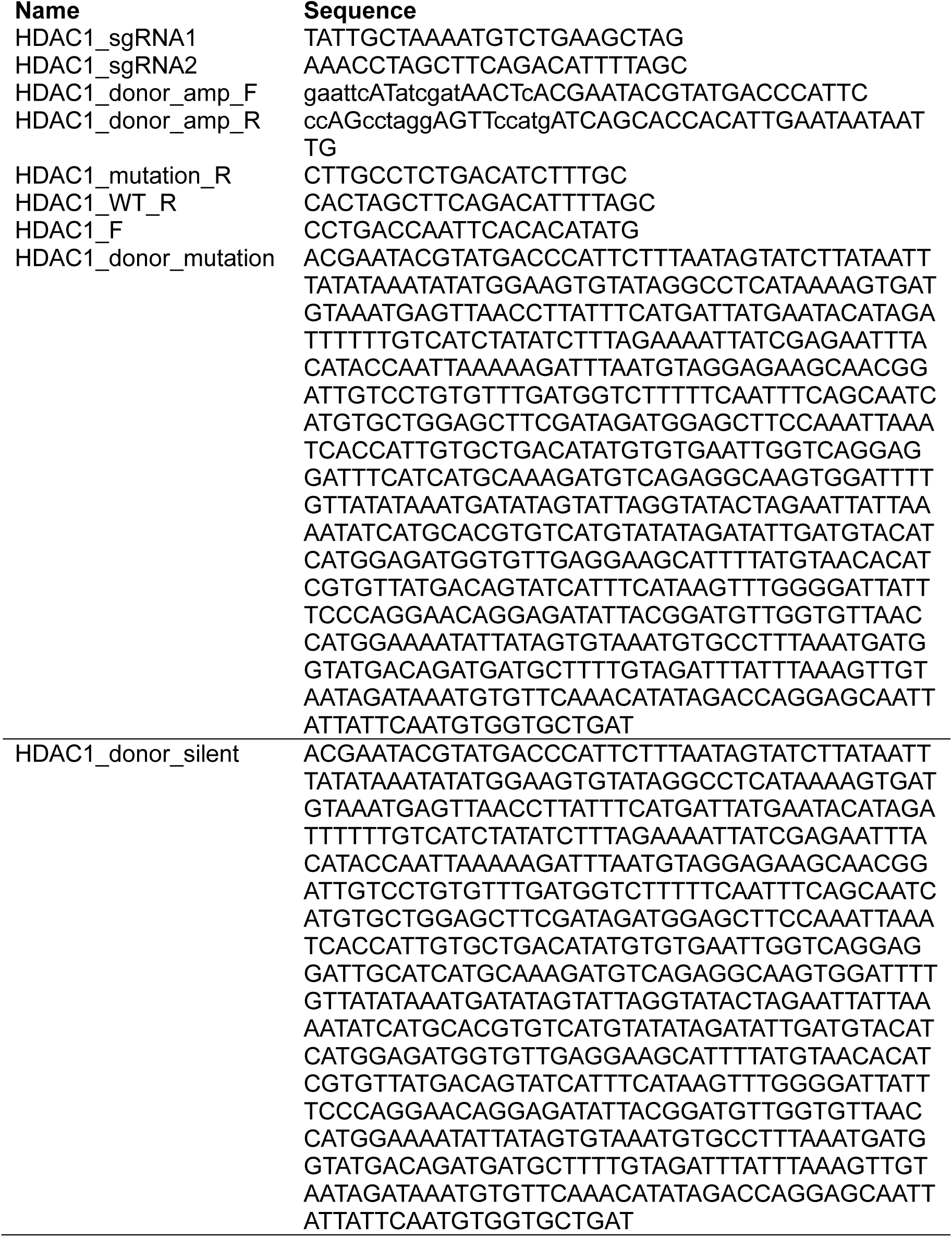
Single guide RNA, donor DNAs, and Sanger sequence primers for plasmid construction and verifying obtained transgenic parasites.

